# Latitudinal MHC variation and haplotype associated differential survival in response to experimental infection of two strains of *Batrachochytrium dendrobatitis* (*Bd*-GPL) in common toads

**DOI:** 10.1101/597559

**Authors:** Maria Cortazar-Chinarro, Sara Meurling, Laurens Schroyens, Mattias Siljestam, Alex Ritcher-Boix, Anssi Laurila, Jacob Höglund

## Abstract

While both innate and adaptive immune system mechanisms have been implicated in resistance against the chytrid fungus *Batrachochytrium dendrobatitis*, studies on the role of specific MHC haplotypes on *Bd* infection are rare. Here, we studied latitudinal variation in MHC Class IIB loci along a latitudinal gradient from southern to northern Sweden in common toads, *Bufo bufo*. Swedish toad populations had fewer MHC Class IIB haplotypes compared to a previous study of populations in Britain. Furthermore, we found MHC diversity to decline from south to the north within Sweden. The low diversity may compromise the ability of northern populations to fight emerging disease, such as the chytrid fungus *Bd*. In a laboratory experiment, we infected newly metamorphosed toads with two strains of the Global Pandemic Lineage of the fungus (*Bd*-GPL) and compared survival with sham controls. We found *Bd*-infected toads had lower survival compared to controls. Survival was dependent on *Bd*-strain and whether experimental toads where collected in the south or the north of Sweden with lower survival in northern individuals. MHC diversity was lower in toads of northern origin, all northern animals being monomorphic for a single MHC haplotype, whereas we found seven different haplotypes in southern animals. Survival of infected animals was dependent on both *Bd*-strain and MHC haplotype suggesting differential infection dynamics depending on both *Bd*-strain and host MHC characteristics.

## Introduction

Emerging infectious diseases are a severe threat to global biodiversity, causing population declines, and species extinctions, across a range of taxa [1]. Amphibian populations world-wide are faced with emerging diseases among them chytrid fungus *Batrachochytrium dendrobatidis* (Bd) which has been shown to infect more than 700 amphibian species and is implicated in the extinction of more than 100 species [2,3]. *Bd* consists of numerous strains and lineages, some of which are highly pathogenic while others are more benign [4, 5]. The infectious lineage *Bd*-GPL has evolved within the last century in south-east Asia and has since spread over the world infecting naïve amphibian species and populations [6]. *Bd*-GPL infections may cause mass mortality, while in other cases populations and species appear resistant to the disease [7]. Studies on the variation in the pathogenicity of the fungus and whether host populations vary in resistance are therefore vital, as is pin-pointing the immunological and genetic basis for such differences. Any amphibian species ability to resist the fungus will vary depending on a range of factors ultimately linked to immunogenetic variation and differences in pathogenicity among different lineages and strains of *Bd* [8].

In general, historical events such as postglacial colonization processes have had a profound impact on the geographical distribution of genetic diversity. During the Last Glacial Maxium (LGM), species inhabiting northern Europe were restricted to refugia in southern Europe ([9], [10]. After the LGM, species expanded northwards, a process which involved repeated founder events [10]. Population size bottle-necks caused by such founder events lead to loss of genetic variation [11]. Therefore populations residing at northern latitudes generally display less genetic diversity as compared to populations residing in former refugia [12,13, 14]. This is due to smaller effective population sizes and more fragmented populations at the edge of the expansion [15, 16].

MHC is a multigene family consisting of various number of genes coding for peptides involved in the adaptive immune defense of vertebrates [17]. It codes for proteins that are part of the adaptive immune system and is associated with disease and parasite resistance [18]. Previous studies of amphibians have shown a direct association between specific MHC alleles and *Bd*-resistance [19, 20].

Understanding geographic variation concerning immuno-genetic diversity is complicated by which mechanisms shape such diversity [21, 22]. For example, the exceptionally high of diversity at the Major Histocompatability Complex (MHC) is believed to be maintained by various forms of balancing selection [23]. Because of the immune response enhancing properties of MHC, it is suggested that heterozygous individuals are more resistant to pathogens as they have a larger available repertoire to recognize pathogens [24]. However, immunogenetic variation may also be shaped by drift which vary in force in and among species [25, 26, 27].

In this study we focused on common toad (*Bufo bufo*) populations in Sweden, an area which was completely glaciated during the LGM. The aims of the study were to quantify MHC Class II variation along a latitudinal gradient, and to study whether *Bd*-mediated mortality depends on the *Bd*-strain, the presence of MHC Class IIB haplotypes or the geographic origin of the experimental animals. We experimentally infected newly metamorphosed common toads from the north and the south of Sweden with two strains of *Bd*-GPL, originating from Sweden and the UK, respectively, or treated toads with a sham control and recorded their survival over 30 days

## Materials and Methods

### MHC variation along the latitudinal gradient

For the analyses of geographical MHC-variation we collected tissue samples from 20 adult toads at 12 locations spread over four different regions along a latitudinal gradient across Sweden (Table 1) in April-May 2015. The toads were caught in breeding aggregations and a small piece of rear leg webbing was removed and immediately stored in 96 % ethanol. DNA was extracted from the tissue by using a DNeasy Blood and Tissue kit (Qiagen) according to the manufacturer’s protocol.

**Table 1.**
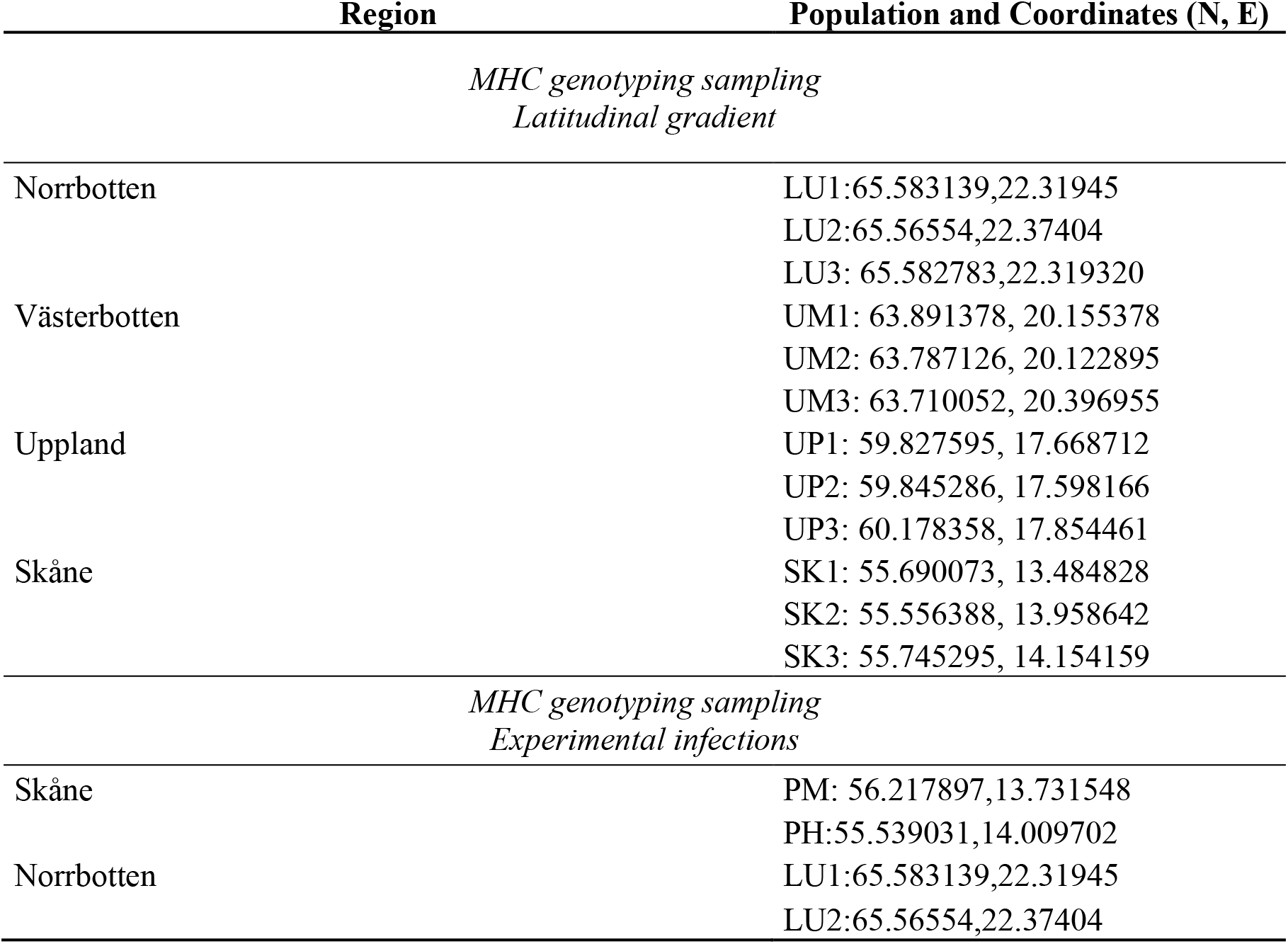
Spatial coordinates of the sampling sites. Tissue from adults from all populations were used in the latitudinal study. For the experiment we used toadlets collected as eggs from LU1 and LU2 (Norrbotten) and PM1 and PH (Skåne). For a map, see Fig. 1.

We amplified MHC Class II exon 2 loci (279 bp) by using the primers for Zeisset [28]. Both forward (2F347F: GTGACCCTCTGCTCTCCATT) and reverse (2R307R: ATAATTCAGTATATACAGGGTCTCACC) primers were modified for Illumina MiSeq sequencing with an individual 8 bp barcode and a sequence of three N to facilitate cluster identification [27], for details. The libraries were constructed by combining 13 different specific barcode tags, four forward primers and nine reverse primers [27].

PCR was done in a 2720 thermal cycler with the protocol: 97°C for 30 sec, followed by 35 cycles of 98°C for 10sec, 57°C for 20 seconds and 72°C for 15sec, and a final round of 72°C for 8 minutes until the samples are cooled down and stored at 4°C. PCR was conducted in a total volume of 30 μL: 0.5μL of DNA and 22.1 μL ddH_2_O, 6 μL Phuside HF buffer, 0.6 μL dNTP, 0.75 μL of both forward and reverse primer and 0.3 μL Phusion Taq. After amplification samples were pre-pooled in five to nine pools in equimolar amounts and purified following the MiniElute Gel Extraction kit (Qiagen). For geographical analyses and for experimental animals eight libraries were prepared with a ThruPlex DNA-Seq Kit (Rubicon genomics) and sequencing was done with Illumina Miseq at NGI Uppsala (Uppsala Genome Center). In total 240 individuals for geography and 160 experimental animals were amplified. Replicates were selected randomly across populations. The Replication rate for the geographic analyses was ~40% (72 samples out of 160). Replication rate for the experimental individuals was 100%.

### Miseq daata analysis

The sequence data was merged using FLASH [29]. The NextGen workbench (www.DnaBaser.com) was used to transform the files to ‘.fasta’ format for further analysis. We demultiplexed the sample sequences with jMHC [30]. An Excel macro [31] was used to estimate allele variants per individual based on Degree of Change (DoC) calculations. The DoC criterion calculates cumulative amplicon sequencing depth among the variants in the amplicon. Plotting the sequencing depth per variant shows an inflection point where the true alleles contribute to a large proportion of the cumulative depth and all the artificial variants only contribute a minor proportion. We assigned the most frequent variants as valid MHC alleles that occurred in at least 3% of the reads [32, 33]. Amplicons with < 300 reads was discarded from analyses for quality reasons. We thoroughly checked for chimeras by eye and any of the valid sequences presented insertions, deletions or stop codons. Verified MHC class IIB exon 2 haplotypes were named in accordance with Klein [34], a species abbreviation followed by gene and a following number e.g. Bubu_DAB*01.

### Experimental infections

Common taod eggs were collected in 2016 at two locations in Skåne (PM and PH, Table 1) and two locations in Luleå (LU1 and LU2). We carefully collected ~ 100 eggs from ~ 10 clutches per pond in order to reduce the risk of sampling related individuals. Eggs were brought to the laboratory in Uppsala and reared in 20 l containers in reconstituted soft water (RSW; NaHCO_3_, CaSO_4_, MgSO_4_ and KCl added to deionized water; APHA 1985). RSW was then used throughout the experiment. Each clutch was kept in a separate container. At metamorphosis (stage 46; Gosner [35] 1960), the tadpoles were removed from the containers and raised until completion of metamorphosis (stage 46; Gosner 1960) in terraria with access to water and shelter. Once the experiments finished, metamorphosed toadlets were euthenized with an overdose of MS222, preserved in 96% ethanol and stored at 4 °C until DNA extraction.

The UK isolate of *Bd* was obtained from the Institute of Zoology, London and isolated from an Alpine newt (*Ichthyosaura alpestris*) from the UK in 2008. The Swedish isolate originated from a Green toad (*Bufotes variabilis*) caught in Norra hamnen, Malmö municipality, Sweden in 2015.

The infection experiment was conducted and animals were housed in the sealed facilities at the Swedish Institute for Veterinary Science, Uppsala. We infected 50 (5 SK1, 14 SK2, 15 LU1, 16 LU2) post metamorphosed toadlets with a *Bd*-GPL strain from the United Kingdom (UK Mal 01, GPL-UK), 52 (4, 17, 16, 15) with an isolate from Sweden (SWED-40-5, GPL-SWED) and 52 (7, 17, 15, 13) with a sham control. Experimental animals were exposed to 12 · 10^6^ zoospores of *Bd* in 30 ml of reconstituted soft water (RSW) (60000 zoospores/ml) for 5 h). Metamorphs in the control group were exposed to an equivalent volume of sterile media and RSW for the same time period, at the same stage of development.

All animals used in the experiment were typed for MHC Class IIB sequence variation according to the same protocol as described above. After experimental infection, the toadlets were kept individually in containers provided with water, shelter and food ad libitum. Survival was recorded daíly and dead animals were preserved in 96% ethanol and stored at 4 °C. Thirty days after infection, the experiment was ended and surviving toadlets were euthanized with an overdose of MS222, preserved in 96% ethanol and stored at 4 °C until DNA extraction.

We used survival until the end of the experiment as a binomial response variable with *Bd*-strain treatment (GPL-UK, GPL-SWED and Control), geographic region / population (depending on the model), presence of various MHC haplotypes as independent variables as well as their interactions in Generalised Linear Models. Due to underdispersion we used a quasibinomial error distribution with the logit link function where the dispersion parameter was estimated to 0.615. Geographic location was tested either as the actual population of origin or classified as region (north or south). Models followed the general structure:

Survival (Yes/No) ~ *Bd*-strain treatment + Region/Population + MHC haplotype + Interactions Model selection was based on Akaike’s Information Criterion (AIC) and all analyses were run in the R packages “car” [36] and “nlme” [37].

## Results

### Geographic MHC variation

At least one MHC Class IIB sequence was successfully amplified in all but one individual. A total of 8 million reads were picked up by the two Miseq runs with on average 2 million reads per library. Thirteen unique sequences (hereafter haplotypes) were identified (Bubu_DAB*1 – Bubu_DAB*13). A maximum of four haplotypes per individual was found in 11 cases, in all other individuals we found one to three haplotypes.

Haplotype number varied between regions and populations (Fig.1) and decreased towards the north. We found significant differences in number of alleles among populations along the latitudinal gradient (F_3,8.010_=10.77, p=0.003; Fig. S1). The observed number of different haplotypes in each region was (from south to north): Skåne 9, Uppland 5, Västerbotten 5 and Norrbotten 4. We found one haplotype unique to Västerbotten (Bubu_DAB*10 found in one individual). In Skåne we found 4 haplotypes unique to this region: Bubu_DAB*9, Bubu_DAB*8, Bubu_DAB*7 and Bubu_DAB*6. Bubu_DAB*01 was found in all individuals in all populations.

**Figure 1.**
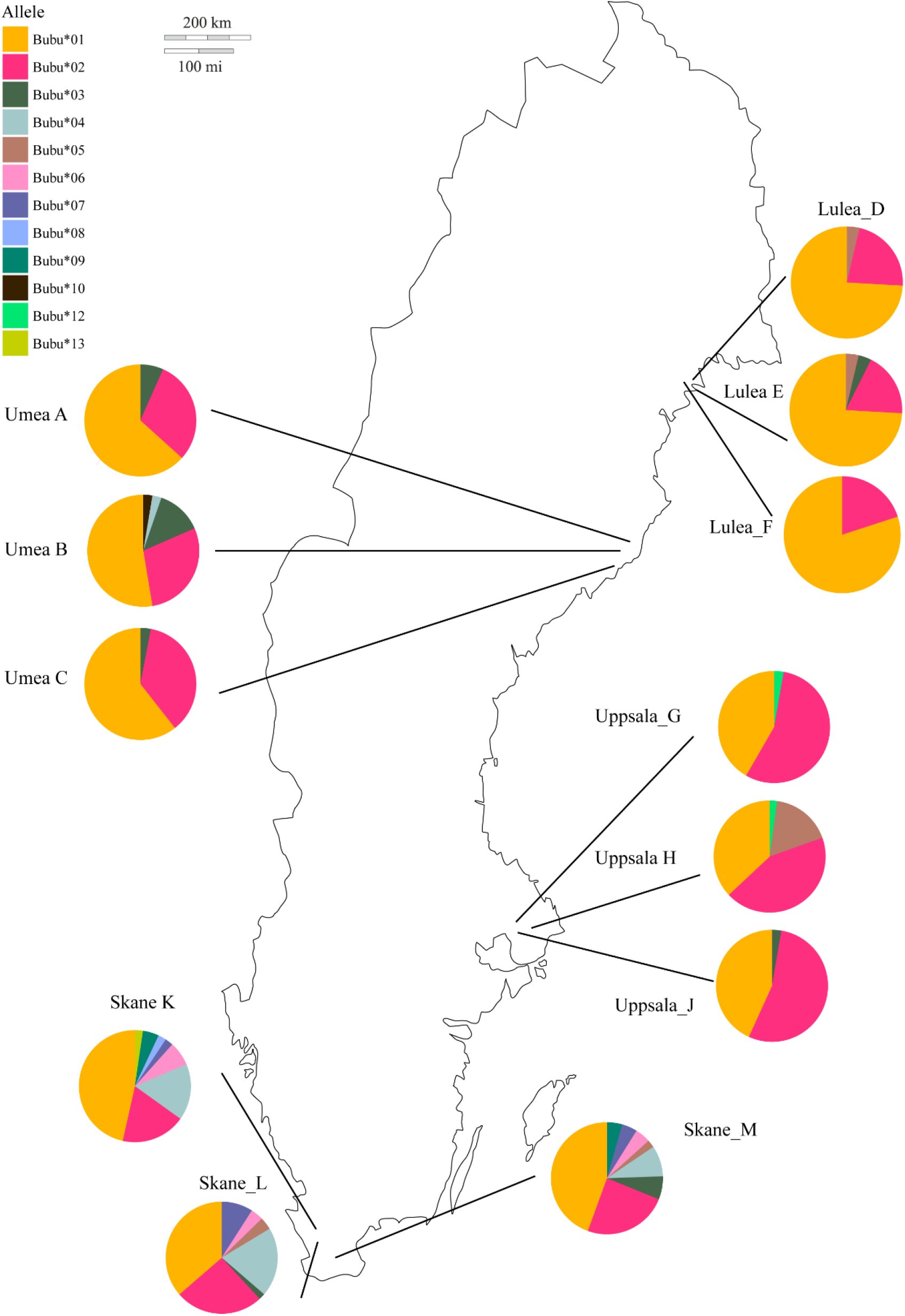
Haplotype frequency distribution of MHC class II sequences in Bufo bufo12 sites in 4 regions in Sweden.

### Experimental infections

Infection was confirmed by qPCR (for details see Meurling et al. in prep). Because of failed reactions infection could not be confirmed or denied in 14.9% of the individuals. One of the exposed individuals (with strain UK) proved to be negative to infection. Low levels of infection could be found in 19.2 *%* of control animals (Table. S1). All animals in the control treatment survived the 30-day experimental period. Of the 102 experimentally infected animals 54 died, 45 from the northern region and nine from the south. Of the six models tested, model 3 including “region” as a main effect and the interactions among “*Bd*-strain” and “MHC haplotype” and “Bd strain” and “region” had the highest AIC followed by a similar model excluding the *“Bd* strain/region” interaction (Table. 2).

**Table 2.**
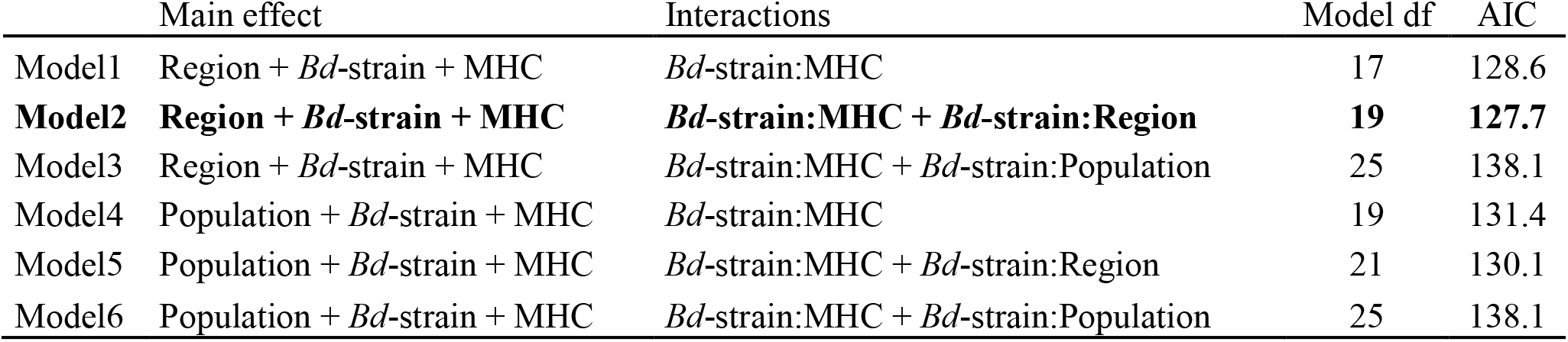
Overview of the different models with main effects and the various interactions modelled. Akaike’s Information Criterion (AIC) was used to select among models.

We discovered a total of seven unique haplotypes: (Bubu_DAB*1, Bubu_DAB*2, Bubu_DAB*4, Bubu_DAB*6, Bubu_DAB*7, Bubu_DAB*9 and Bubu_DAB*11) in the experimental animals. All seven were found in the southern region, whereas in the northern region all individuals were monomorphic for Bubu_DAB*1.

Of the six models tested, model 2 was chosen having the lowest AIC (Table. 2). The model includes “Region”, “*Bd*-strain”, and “MHC-haplotype” as a main effects and the interactions among “*Bd*-strain” and “MHC haplotype” as well as *“Bd* strain” and “Region” (Table. 2). The result from the model are presented in table 3.

**Table 3.**
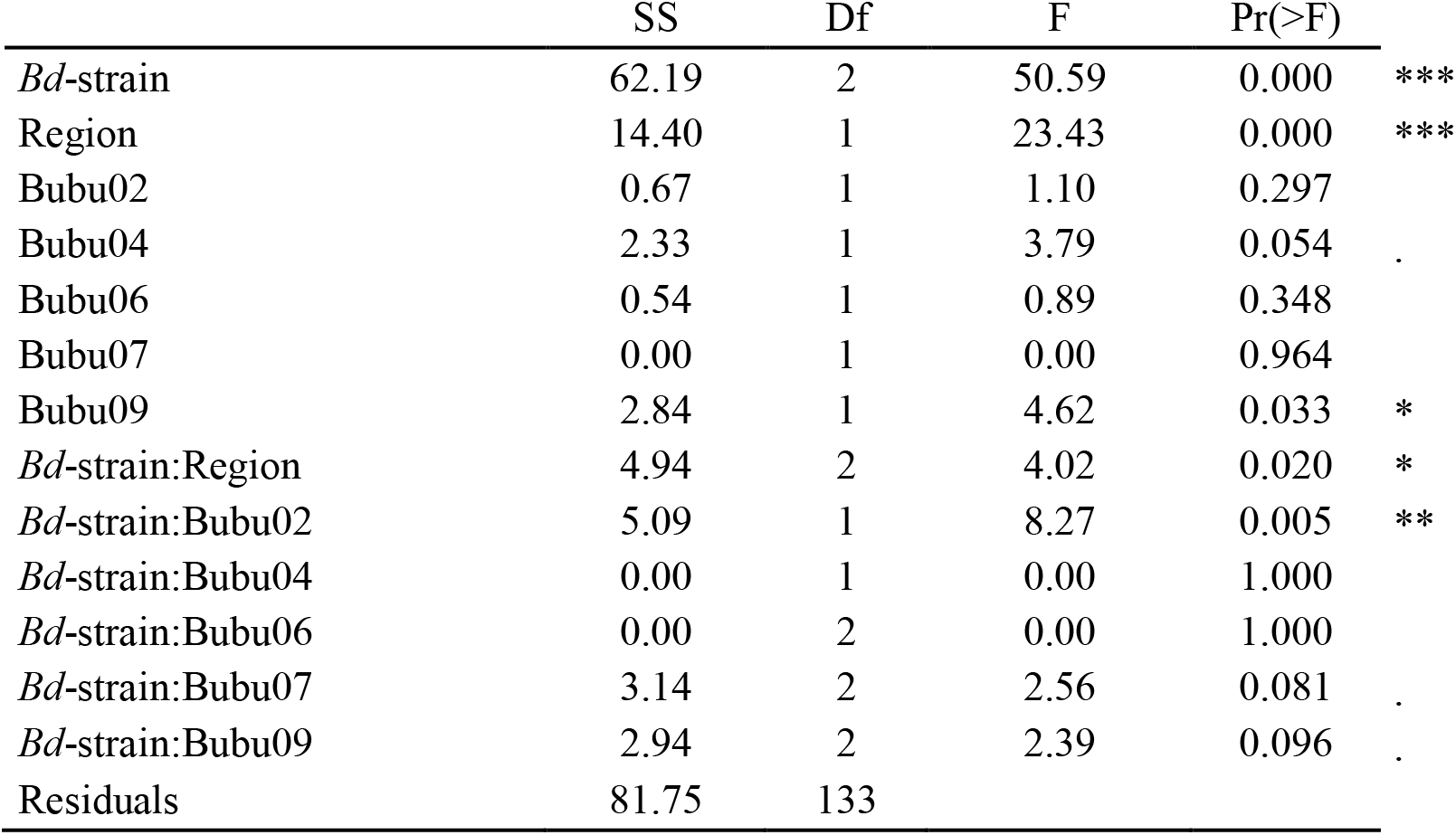
Results of GLM in the best model (model 2).

Of the interactions in the best model, a significant interaction among “*Bd*-strain” and “region” indicate that increased survival in the south depends on which *Bd*-strain the toads were infected with (p=0.020; F=4.02; Fig. 2). Furthermore, the haplotype Bubu_DAB*2 appears to provide protection against the Swedish *Bd* strain but this haplotype was detrimental when toads were infected with the UK-strain (p=0.005; F=8.27; Fig. 3a). The haplotype Bubu_DAB*9 had a detrimental effect on survival with a possible interaction with the *Bd*-strain treatment being marginally less detrimental for individuals infected by the Swedish strain (p=0.096; F=2.39; Fig. 3b).

**Figure 2.**
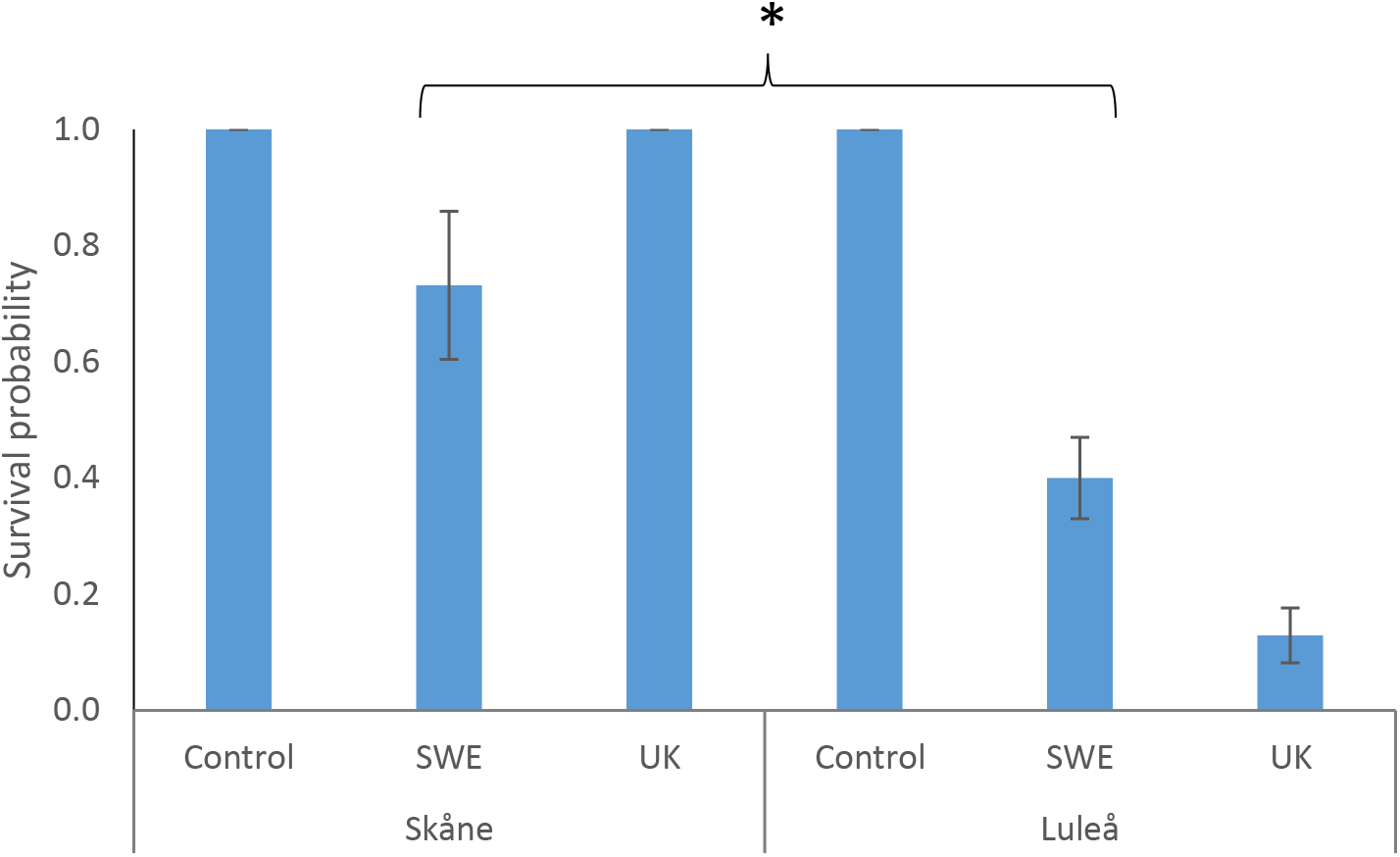
Toads only carrying Bubu_DAB*1 showed an increased survival in the South compared to the North, but the increase depends on *Bd*-strain. Hence, the survival probability assumes that none of the MHC-alleles considered in the model are carried.

**Figure 3.**
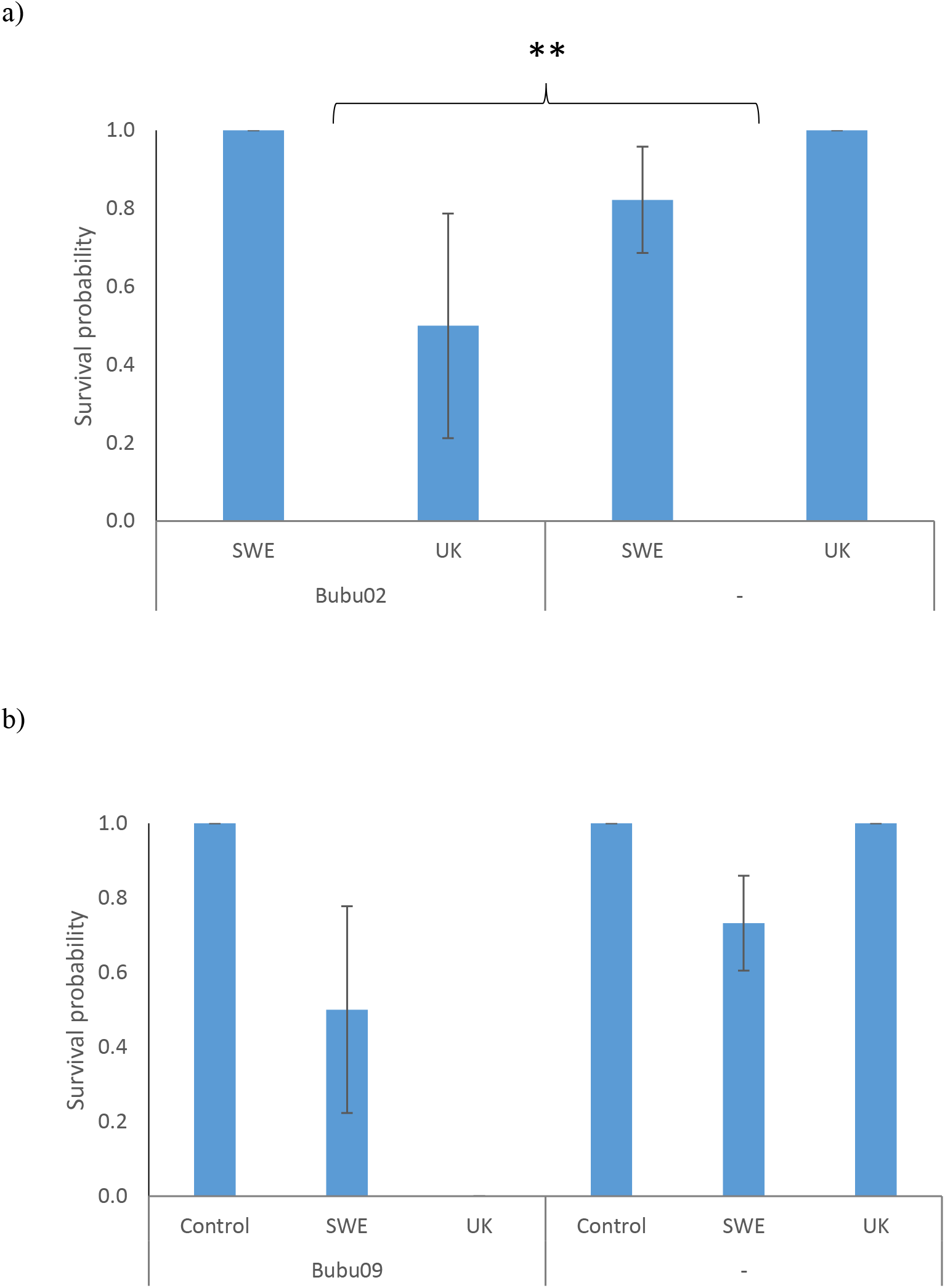
The predicted effect of the three alleles with potential *Bd*-strain interactions. a) The Bubu_DAB*2 haplotype provided protection against the Swedish *Bd*-strain, but was detrimental for individuals infected with the UK *Bd*-strain. The predicted values assume using a background of carrying the Bubu_DAB*6 and Bubu_DAB*7 haplotype as this was the case for all three individuals carrying the Bubu_DAB*2 haplotype. None of the three individuals with Bubu_DAB*2 were present in the control treatment, which is thereby excluded. b) Bubu_DAB*9 haplotype has a detrimental effect on survival for *Bd*-infected individuals, especially for the UK-strain. All predicted values assume the region south as this was the case for all individuals carrying these MHC-alleles.

## Discussion

Previous studies of latitudinal MHC variation in amphibians suggest neutral processes have had a strong influence on the geographic distribution of MHC diversity over large biogeographical areas. Thus MHC diversity tends to decline at northern latitudes when moving away from putative post glacial refugia in a number of species [38, 32, 39, 40, 27, 41]. A study of *B. bufo* MHC Class IIB variation in the UK found 17 unique sequences in 149 animals [28], and we found a total of 13 in 239 animals further to the north in Sweden, three sequences of which were also found in the UK. We also found lower MHC diversity at higher latitudes in Sweden.

Regional variation in MHC diversity may be shaped by both genetic drift and selection, and populations at different latitudes may face varying strengths and forms of selection [27, 42]. Varying selection may depend on differences in parasites and pathogen pressure [43], but also on historical demographic process such as population size fluctuations mediated through bottlenecks and founder events [11]. Generally, amphibian populations tend to be smaller and more fragmented at northern latitudes [44, 15], and since the force of genetic drift is inversely related to population size [45], northern populations are generally predicted to be more influenced by drift. Conversely in southern populations, selection is expected to be relatively more influential as populations tend to be larger and more connected. This has consequences for population’s ability to combat disease and pathogens since immunogenetic diversity is predicted to be relatively less shaped by selection at northern latitudes [42].

In the experimental animals, we observed higher mortality in northern individuals. Whether this result depends on the observed lack of MHC variation or some other factor which differs among the northern and southern regions is impossible to discern with this study. Interestingly, we found a difference between the regions in response to the two different *Bd*-strains. *Bd* had a negative impact on survival regardless of strain but when infected with the UK strain, the southern individuals had almost complete survival whereas the northern ones suffered high mortality. When infected with the Swedish strain, both southern and northern individuals had high mortality albeit the rate was higher in the north. Unfortunately, due to lack of MHC variation in the northern animals it is impossible to discern whether this was due to differences in MHC variation or some other factor.

However, we were able to pick up some MHC-related differences in survival as presence of the haplotype Bubu_DAB*9 seemed to have a negative impact on survival when infected with the UK-strain, but the effect was limited when infected with the Swedish strain. Similarly, the haplotype Bubu_DAB*2 had a negative impact on survival when infected with the UK-strain but was marginally beneficial when infected by the Swedish. This difference among strains is somewhat surprising as both strains used in the experiment belong to *Bd*-GPL lineage which has emerged within the last century and is the cause of the extinction of more than 100 amphibian species [6]. Non GPL *Bd*-strains are rarely associated with chytridiomycosis, population declines and species extinction [6].

To our knowledge, this is the first report of differences in host mortality among *Bd*-GPL strains. Previous studies have shown MHC Class IIB-haplotype related differences in survival in *Lithobates yavapaiensis* [19, 20]. In *Litoria verrauxii*, experimental studies showed that certain MHC Class IIB amino acid sequences conferred a survival advantage and the differences were ascribed to variation at the peptide-binding coding residues of the amino acid sequences [46]. The haplotypes which conferred a survival advantage in *L. verrauxii* were then compared to the most common haplotypes observed in species classified as either susceptible or resistant to *Bd* and *B. bufo* was grouped among the resistant species [46]. However, *B. bufo* have been shown to suffer high mortality in experimental infections with *Bd* [47]. Also, *Bd* has been shown to infect the sister species *B. spinosus* in a Spanish population (classified as *B. bufo* at the time of the study). While *Bd* was associated with mortality, no population declines were reported [48]. Yet, the species has declined in the study area since the time of the study [49].

The available evidence thus strongly suggests that susceptibility to *Bd* depends on a range of factors which may explain both intra and interspecific differences. Intraspecific differences may be ascribed to immuno-genetic competence and diversity. Populations which have a long history of exposure to *Bd* may have undergone selection favouring resistance alleles. Populations naïve to the disease may by chance have resistance alleles but populations further away from refugia and core areas of the distribution are often depauperate in genetic variation and less likely to harbour adaptive variation. Similar processes may act at the interspecific level as different species are shaped by different demographic events. For example, the moor frog in northern Europe harbours quite extensive MHC Class IIB variation, while the natterjack toad, *Epidalea calamita*, have lost most of its MHC-variation during the postglacial expansion to northern Europe [40, 50]. North American ranid frogs have been shown to show low hazard ratios while bufonids showed high when exposed to *Bd* [7]. The different strains of the fungus may also vary in susceptibility within and among populations. *B. dendodrabatidis* and *B. salamandrivorans* have been shown to display quite different infection strategies, not surprisingly considering the deep divergence among these two chytrids [51]. As indicated by this study, also recently diverged *Bd*-strains may vary in susceptibility although this is unlikely to be dependent on vastly different infection strategies. Nevertheless, the constant coevolutionary arms-race among hosts and pathogens is likely to lead to a range of susceptibilities and pathogenicity within and among species [52,53].

## References

[1] Daszak, P., Cunningham, A.A. & Hyatt, A.D. 2000 Wildlife ecology - Emerging infectious diseases of wildlife - Threats to biodiversity and human health. Science 287, 443–449. (doi: 10.1126/science.287.5452.443).

[2] Fisher, M.C., Garner, T.W. & Walker, S.F. 2009 Global emergence of Batrachochytrium dendrobatidis and amphibian chytridiomycosis in space, time, and host. Annual review of microbiology 63, 291–310.

[3] Lips, K.R. 2016 Overview of chytrid emergence and impacts on amphibians. Philosophical Transactions of the Royal Society B: Biological Sciences 371, 20150465.

[4] Refsnider, J.M., Poorten, T.J., Langhammer, P.F., Burrowes, P.A. & Rosenblum, E.B. 2015 Genomic correlates of virulence attenuation in the deadly amphibian chytrid fungus, Batrachochytrium dendrobatidis. G3: Genes, Genomes, Genetics 5, 2291–2298.

[5] Dang, T.D., Searle, C.L. & Blaustein, A.R. 2017 Virulence variation among strains of the emerging infectious fungus Batrachochytrium dendrobatidis (Bd) in multiple amphibian host species. Diseases of aquatic organisms 124, 233–239.

[6] O’Hanlon, S.J., Rieux, A., Farrer, R.A., Rosa, G.M., Waldman, B., Bataille, A., Kosch, T.A., Murray, K.A., Brankovics, B. & Fumagalli, M. 2018 Recent Asian origin of chytrid fungi causing global amphibian declines. Science 360, 621–627.

[7] Gervasi, S.S., Stephens, P.R., Hua, J., Searle, C.L., Xie, G.Y., Urbina, J., Olson, D.H., Bancroft, B.A., Weis, V. & Hammond, J.I. 2017 Linking ecology and epidemiology to understand predictors of multi-host responses to an emerging pathogen, the amphibian chytrid fungus. PLoS One 12, e0167882.

[8] Grogan, L.F., Robert, J., Berger, L., Skerratt, L.F., Scheele, B.C., Castley, J.G., Newell, D.A. & McCallum, H.I. 2018 Review of the amphibian immune response to chytridiomycosis, and future directions. Frontiers in Immunology 9, 2536.

[9] Hewitt, G.M. 2004 The structure of biodiversity - insights from molecular phylogeography. Front. Zool. 1, 4–4.

[10] Hampe, A. & Petit, R.J. 2005 Conserving biodiversity under climate change: the rear edge matters. Ecol. Lett. 8, 461–467. (doi:10.1111/j.1461-0248.2005.00739.x).

[11] Nei, M., Maruyama, T. & Chakraborty, R. 1975 Bottleneck effect and genetic-variability in populations. Evolution 29, 1–10. (doi:10.2307/2407137).

[12] Hewitt, G.M. 1996 Some genetic consequences of ice ages, and their role in divergence and speciation. Biol. J. Linn. Soc. 58, 247–276. (doi: 10.1006/bijl.1996.0035).

[13] Hewitt, G.M. 1999 Post-glacial re-colonization of European biota. Biol. J. Linn. Soc. 68, 87–112. (doi: 10.1006/bijl.1999.0332).

[14] Taberlet, P., Fumagalli, L., Wust-Saucy, A.G. & Cosson, J.F. 1998 Comparative phylogeography and postglacial colonization routes in Europe. Mol. Ecol. 7, 453–464.

[15] Eckert, C.G., Samis, K.E. & Lougheed, S.C. 2008 Genetic variation across species’ geographical ranges: the central-marginal hypothesis and beyond. Mol. Ecol. 17, 1170–1188. (doi:10.1111/j.1365-294X.2007.03659.x).

[16] Guo, Q. 2012 Incorporating latitudinal and central–marginal trends in assessing genetic variation across species ranges. Mol. Ecol. 21, 5396–5403.

[17] Janeway, C.A., Travers, P., Walport, M. & Shlomchik, M. 2001 Immunobiology: the immune system in health and disease, Garland Pub. New York.

[18] Hedrick, P.W. 1994 EVOLUTIONARY GENETICS OF THE MAJOR HISTOCOMPATIBILITY COMPLEX. Am. Nat. 143, 945–964. (doi:10.1086/285643).

[19] Savage, A.E. & Zamudio, K.R. 2011 MHC genotypes associate with resistance to a frog-killing fungus. Proc. Natl. Acad. Sci. U. S. A. 108, 16705–16710. (doi:10.1073/pnas.1106893108).

[20] Savage, A.E. & Zamudio, K.R. 2016 Adaptive tolerance to a pathogenic fungus drives major histocompatibility complex evolution in natural amphibian populations. Proc. R. Soc. Lond. B. Biol. Sci. 283. (doi:10.1098/rspb.2015.3115).

[21] Potts, W.K. & Slev, P.R. 1995 Pathogen-based models favoring MHC genetic diversity. Immunol. Rev. 143, 181–197.

[22] Hedrick, P.W. 1999 Perspective: highly variable loci and their interpretation in evolution and conservation. Evolution 53, 313–318.

[23] Hughes, A.L. & Nei, M. 1988 Pattern of nucleotide substitution at major histocompatibility complex class I loci reveals overdominant selection. Nature 335, 167.

[24] Doherty, P.C. & Zinkernagel, R.M. 1975 Enhanced immunological surveillance in mice haterozygous at H-2 gene complex. Nature 256, 50–52.

[25] Piertney, S.B. & Oliver, M.K. 2006 The evolutionary ecology of the major histocompatibility complex. Heredity 96, 7–21. (doi:10.1038/sj.hdy.6800724).

[26] Spurgin, L.G. & Richardson, D.S. 2010 How pathogens drive genetic diversity: MHC, mechanisms and misunderstandings. Proc. R. Soc. Lond. B. Biol. Sci. 277, 979–988. (doi:10.1098/rspb.2009.2084).

[27] Cortázar-Chinarro, M., Lattenkamp, E.Z., Meyer-Lucht, Y., Luquet, E., Laurila, A. & Höglund, J. 2017 Drift, selection, or migration? Processes affecting genetic differentiation and variation along a latitudinal gradient in an amphibian. BMC Evol. Biol. 17, 189.

[28] Zeisset, I. & Beebee, T.J.C. 2013 Bufo MHC class II loci with conserved introns flanking exon 2: cross-species amplification with common primers. Conserv. Genet. Resour. 5, 211–213. (doi: 10.1007/s12686-012-9770-y).

[29] Magoč, T. & Salzberg, S.L. 2011 FLASH: fast length adjustment of short reads to improve genome assemblies. Bioinformatics 27, 2957–2963.

[30] Stuglik, M.T., Radwan, J. & Babik, W. 2011 jMHC: software assistant for multilocus genotyping of gene families using next-generation amplicon sequencing. Mol. Ecol. Resour. 11, 739–742. (doi: 10.1111/j.1755-0998.2011.02997.x).

[31] Lighten, J., Van Oosterhout, C. & Bentzen, P. 2014 Critical review of NGS analyses for de novo genotyping multigene families. Mol. Ecol. 23, 3957–3972. (doi:10.1111/mec.12843).

[32] Babik, W., Pabijan, M., Arntzen, J.W., Cogalniceanu, D., Durka, W. & Radwan, J. 2009 Long-term survival of a urodele amphibian despite depleted major histocompatibility complex variation. Mol. Ecol. 18, 769–781. (doi:10.1111/j.1365-294X.2008.04057.x).

[33] Galan, M., Guivier, E., Caraux, G., Charbonnel, N. & Cosson, J.-F. 2010 A 454 multiplex sequencing method for rapid and reliable genotyping of highly polymorphic genes in large-scale studies. BMC Genomics 11. (doi:10.1186/1471-2164-11-296).

[34] Klein, J. 1975 Biology of the mouse histocompatibility-2 complex. Principles of immunogenetics applied to a single system, Springer-Verlag.

[35] Gosner, K.L. 1960 A simplified table for staging anuran embryos and larvae with notes on identification. Herpetologica 16, 183–190.

[36] Fox, J., Weisberg, S., Adler, D., Bates, D., Baud-Bovy, G., Ellison, S., Firth, D., Friendly, M., Gorjanc, G. & Graves, S. 2012 Package ‘car’. Vienna: R Foundation for Statistical Computing.

[37] Pinheiro, J., Bates, D., DebRoy, S., Sarkar, D., Heisterkamp, S., Van Willigen, B. & Maintainer, R. 2017 Package ‘nlme’. Linear and Nonlinear Mixed Effects Models, version, 3–1.

[38] Babik, W., Pabijan, M. & Radwan, J. 2008 Contrasting patterns of variation in MHC loci in the Alpine newt. Mol. Ecol. 17, 2339–2355.

[39] Wielstra, B., Babik, W. & Arntzen, J.W. 2015 The crested newt Triturus cristatus recolonized temperate Eurasia from an extra-Mediterranean glacial refugium. Biol. J. Linn. Soc. 114, 574–587.

[40] Zeisset, I. & Beebee, T.J.C. 2014 Drift Rather than Selection Dominates MHC Class II Allelic Diversity Patterns at the Biogeographical Range Scale in Natterjack Toads Bufo calamita. PLoS One 9. (doi:e10017610.1371/journal.pone.0100176).

[41] Talarico, L., Babik, W., Marta, S. & Mattoccia, M. 2019 Genetic drift shaped MHC IIB diversity of an endangered anuran species within the Italian glacial refugium. Journal of Zoology.

[42] Cortazar-Chinarro, M., Meyer-Lucht, Y., Laurila, A. & Höglund, J. 2018 Signatures of historical selection on MHC reveal different selection patterns in the moor frog (Rana arvalis). Immunogenetics 70, 477–484.

[43] Sommer, S. 2005 The importance of immune gene variability (MHC) in evolutionary ecology and conservation. Front. Zool. 2, 1–18.

[44] Johansson, M., Primmer, C.R. & Merila, J. 2006 History vs. current demography: explaining the genetic population structure of the common frog (Rana temporaria). Mol. Ecol. 15, 975–983. (doi: 10.1111/j.1365-294X.2006.02866.x).

[45] Wright, S. 1931 Evolution in Mendelian populations. Genetics 16, 0097–0159.

[46] Bataille, A., Cashins, S.D., Grogan, L., Skerratt, L.F., Hunter, D., McFadden, M., Scheele, B., Brannelly, L.A., Macris, A., Harlow, P.S., et al. 2015 Susceptibility of amphibians to chytridiomycosis is associated with MHC class II conformation. Proc. R. Soc. Lond. B. Biol. Sci. 282. (doi: 10.1098/rspb.2014.3127).

[47] Garner, T., Garcia, G., Carroll, B. & Fisher, M. 2009 Using itraconazole to clear Batrachochytrium dendrobatidis infection, and subsequent depigmentation of Alytes muletensis tadpoles. Diseases of aquatic organisms 83, 257–260.

[48] Bosch, J. & Martínez-Solano, I. 2006 Chytrid fungus infection related to unusual mortalities of Salamandra salamandra and Bufo bufo in the Penalara Natural Park, Spain. Oryx 40, 84–89.

[49] Bosch, J., Fernández-Beaskoetxea, S., Garner, T.W. & Carrascal, L.M. 2018 Long-term monitoring of an amphibian community after a climate change-and infectious disease-driven species extirpation. Global change biology 24, 2622–2632.

[50] Hoglund, J., Wengstrom, A., Rogell, B. & Meyer-Lucht, Y. 2015 Low MHC variation in isolated island populations of the Natterjack toad (Bufo calamita). Conserv. Genet. 16, 1007–1010. (doi: 10.1007/s10592-015-07151-3).

[51] Farrer, R.A., Martel, A., Verbrugghe, E., Abouelleil, A., Ducatelle, R., Longcore, J.E., James, T.Y., Pasmans, F., Fisher, M.C. & Cuomo, C.A. 2017 Genomic innovations linked to infection strategies across emerging pathogenic chytrid fungi. Nat. Commun. 8, 11. (doi:10.1038/ncomms14742).

[52] L, V.V. 1973 A new evolutionary law. Evol Theor 1, 1–30.

[53] Ejsmond, M.J. & Radwan, J. 2015 Red Queen Processes Drive Positive Selection on Major Histocompatibility Complex (MHC) Genes. PLoS Comput. Biol. 11. (doi: 10.1371/journal.pcbi.1004627).

