## supplemental Figure 1 and Table 1 for "Latitudinal MHC variation and haplotype associated differential survival in response to experimental infection of two strains of *Batrachochytrium dendrobatitis* (*Bd*-GPL) in common toads"

**Suplementary Material for:**

**This Pdf file includes:**

**Figure S1. Number of MHC class II allelic variants per individual.**

**Table S1. Summary of the infection individuals in the two *Bd*-treatments.**

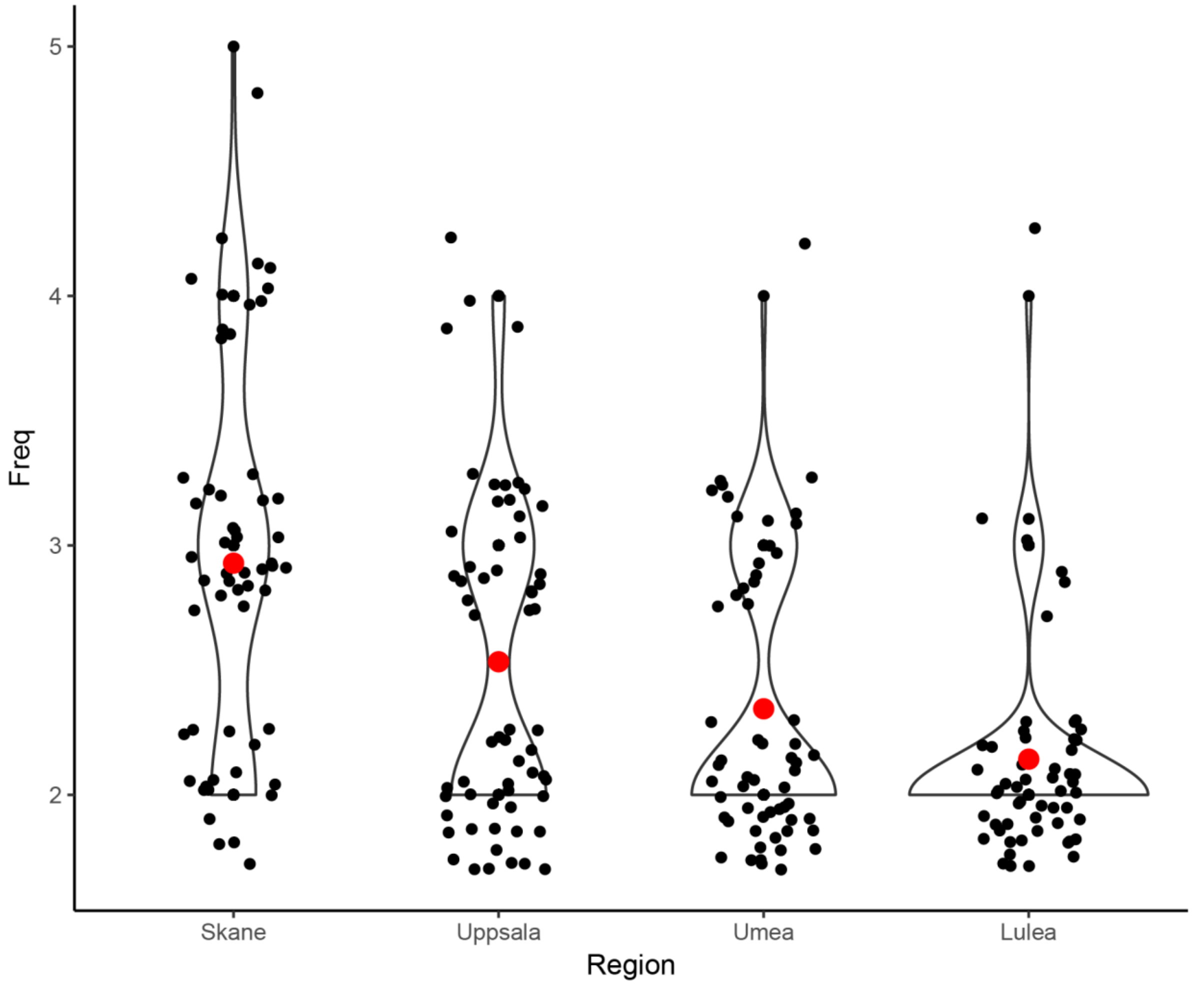


**Fig S1. Number of MHC class II allelic variants per individual.** Every black circle corresponds to one individuals and the red circle represents the mean number of MHC class II variants present in every Region south to north following the latitudinal gradient, in this order: Skåne, Uppsala, Umeå and Luleå.

**Table S1. Summary of the infection individuals in the two Bd-treatments.** Number of positive, negative and the Infection rate confirmed by qPCR in individuals infected with the GPL-UK, GPL-SWE Bd-strains and the control.

| | Positive | % | Negative | % | Failed | % | Total |
|---|---|---|---|---|---|---|---|
| **UK** | 45 | 90.000 | 1 | 2.000 | 4 | 8.000 | 50 |
| **SWE** | 46 | 88.462 | 0 | 0.000 | 6 | 11.538 | 52 |
| **Control** | 10 | 19.231 | 29 | 55.769 | 13 | 25.000 | 52 |
| **Total** | **101** | **197.692** | **30** | **57.769** | **23** | **44.538** | **154** |
